# Enzyme stability-activity trade-off : new insights from protein stability weaknesses and evolutionary conservation

**DOI:** 10.1101/2023.05.02.539073

**Authors:** Qingzhen Hou, Marianne Rooman, Fabrizio Pucci

## Abstract

A general limitation of the use of enzymes in biotechnological processes under sometimes non-physiological conditions is the complex interplay between two key quantities, enzyme activity and stability, where the increase of one is often associated with the decrease of the other. A precise stability-activity trade-off is necessary for the enzymes to be fully functional, but its weight in different protein regions and its dependence on environmental conditions is not yet elucidated. To advance this issue, we used the formalism that we have recently developed to effectively identify stability strength and weakness regions in protein structures, and applied it to a large set of globular enzymes with known experimental structure and catalytic sites. Our analysis showed a striking oscillatory pattern of free energy compensation centered on the catalytic region. Indeed, catalytic residues are usually non-optimal with respect to stability, but residues in the first shell around the catalytic site are, on the average, stability strengths and thus compensate for this lack of stability; residues in the second shell are weaker again, and so on. This trend is consistent across all enzyme families. It is accompanied by a similar, but less pronounced, pattern of residue conservation across evolution. In addition, we analyzed cold- and heat-adapted enzymes separately and highlighted different patterns of stability strengths and weaknesses, which provide insight into the longstanding problem of catalytic rate enhancement in cold environments. The successful comparison of our stability and conservation results with experimental fitness data, obtained by deep mutagenesis scanning, led us to propose criteria for improving catalytic activity while maintaining enzyme stability, a key goal in enzyme design.

## 1 Introduction

Enzymes are widely used as efficient biological catalysts in an large series of biotechnological and biopharmaceutical applications [5]. In the last decades, a lot of studies have been devoted to design new enzymes with improved stability and turnover by computational and/or experimental approaches [8, 43, 26, 11, 42, 27, 9, 22]. Despite these valuable contributions, it is still unclear how the biophysical principles have shaped the structural stability and dynamics of the catalytic regions of enzymes, which in turn determine their characteristic high efficiency and substrate specificity.

Enzyme catalytic sites are further constrained by the environment in which they must be fully functional since their host organisms have adapted to sometimes extreme environmental conditions in terms of, e.g., pH, temperature and ion concentration [34, 1, 39, 12, 3]. For example, some enzymes have evolved to achieve high activity at low temperatures by tuning the flexibility of their catalytic regions [36, 37], even though recent computational approaches have instead suggested that mostly certain regions outside of the active site boost entropic contributions to achieve enzymatic rate enhancement [2, 33].

More generally, the stability-function trade-off of enzymes leads to catalytic regions that are clearly not optimized for stability. They correspond to stability weaknesses in the language we introduced earlier [19, 13] or, almost equivalently, to highly frustrated regions [18]. While the key catalytic residues usually remain intact during natural evolution, mutations that modulate the affinity and the turnover rates for substrates are often localized in the periphery of the active site [38]. However, the complex interaction between the stability strengths and weaknesses of the catalytic residues, their periphery and the rest of the enzymes remains elusive because no experimental method can have direct access to this information. Elucidating this could be of great help in designing new enzymes with increased activity and specificity.

To advance this issue, we used a computational approach based on the well-known formalism of statistical potentials [15] to determine strengths and weakness regions [19, 13] in enzymes. This method has proven to be accurate but also much faster than standard molecular dynamics simulations and can therefore be applied for proteome-scale analyses. By doing so, we have better understood the stability-function trade-off of enzymes, but also how evolution and environmental constraints act to tune this interplay and improve catalytic properties.

## Methods

### Dataset of enzyme structures

We set up a dataset of globular enzymes with annotated catalytic residues and mechanism. For this purpose, we started by downloading the Catalytic Site Atlas (CSA) [29], and identified the list of enzymes with known catalytic properties of which the experimental 3-dimensional (3D) structure is available in the Protein DataBank (PDB) [7]. From this initial set, we removed the structures that have mutations or post-translational modifications in the catalytic site. This resulted in a set of 883 enzymes.

We then used the software PISCES [40] to select the subset of structures that were determined by X-ray crystallography with a resolution of 2.5 Å at most and have a maximum sequence identity of 25% with any other entry of the dataset. We thus obtained a set of 633 enzyme structures. Importantly, we considered the biologically active quaternary structure, and therefore selected the structures of the biological units (biounits) assigned by the authors of the submission in the PDB; if this information is unavailable, we took the biounit predicted by the PISA program [23].

We applied a last filter to the enzyme dataset by limiting our analysis to globular (non-membrane) proteins and dropping hetero-oligomers as well as large homo-oligomeric structures containing more than eight chains. The final dataset of enzyme structures, referred to as 𝒟_*CSA*_, consists of 551 entries among which 240 monomers and 311 homooligomers. The list of all enzymes and their PDB structure are available in our repository github.com/3BioCompBio/EnzymeStability.

A crucial parameter in our analyses is the spatial distance of a residue *i* in a given chain of an enzyme to the active site. We defined this distance as the closest distance between any heavy atom of residue *i* and any heavy atom of the catalytic residues in all the chains of the enzyme. We chose this distance definition to avoid as much as possible the impact of the size and geometry of the catalytic site. Indeed, considering instead the distance to the geometric center of the catalytic site would mix up, e.g., residues that are close to a large catalytic site and residues that are distant from a small catalytic site.

### Identification of stability strength and weakness regions

In order to identify stability strengths and weaknesses in a protein structure, we used the SWOTein program that we developed earlier [15, 13]. We briefly review the main ideas behind SWOTein and refer the reader to Supplementary Section S1 and to the original papers for details.

Statistical potentials are one of the key ingredients of the SWOTein algorithm. They are knowledge-based mean force potentials derived from frequencies of sequence-structure associations computed from a well curated dataset of experimentally resolved protein structures. More precisely, the free energy of the association between a given structure element *c* and a sequence element *s* is obtained from the frequency of observation of their association (*c, s*) in the structure dataset using the inverse Boltzmann law.

SWOTein [15] uses three different kinds of statistical potentials, noted *acc, tor* and *dis*. In these potentials, the *c* element is either the solvent accessibility of a given residue (*acc*), its backbone torsion angle domain (*tor*) or the inter-residue distance between two residues (*dis*), and the sequence element *s* is a pair of amino acid types. Using these potentials and partitioning the free energy into per-residue contributions, we defined three folding free energy values for each residue *i*, 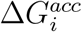, 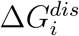 and 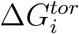, which we linearly combined into a unique per-residue folding free energy contribution as:

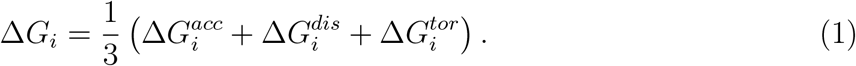

Note that we just summed the free energy contributions without adding weight factors. We made this choice because we prefer avoiding additional parameters and because there are no experimental measurements of the folding free energy per residue that can be used to identify these parameters.

With our conventions, negative Δ*G*_*i*_ values identify regions called stability strengths, which strongly contribute to the stability of the overall protein structure. Positive Δ*G*_*i*_ values rather indicate stability weaknesses, which correspond to regions that are not stable by themselves but are often optimized for functional reasons rather than for their contribution to the stability of the native structure. [19, 4]

## 2 Results

Investigating the biophysical and structural features of catalytic regions in enzymes is crucial to understand their role in the reactions they catalyze, and this understanding is necessary to be able to rationally modify enzyme specificity and turnover. Here we performed a systematic analysis of the patterns of stability and evolutionary conservation inside and outside catalytic regions in a large dataset of enzyme structures. We also examined whether these patterns differ between cold- or heat-adapted enzymes, and whether they correlate with experimentally measured fitness.

### Strengths and weaknesses in catalytic regions

We first analyzed the average pattern of stability strengths and weaknesses in catalytic sites and their periphery. For this purpose, we considered three types of statistical potentials labeled *dis, acc*, and *tor*, which are based on the preferences of amino acids to be separated by a certain distance, to have a certain solvent accessibility, or to adopt a certain backbone torsion angle (see Methods). Using these three potentials, we computed the folding free energy contributions 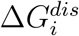, 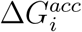 and 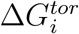 of every residue *i* of the enzyme dataset 𝒟_*CSA*_ as well as Δ*G*_*i*_, the combination of these contributions defined in Eq. (1). In parallel, we computed the distance *d*_*i*_ of each residue *i* to the closest catalytic residue (see Methods).

The per-residue folding free energy contributions as a function of the distance to the active site, averaged over all proteins of the 𝒟_*CSA*_ set are shown in Fig. 1. Examples of per-protein curves are given in Fig. 2 and Supplementary Fig. S1 for four enzymes; the plots for every enzyme from 𝒟_*CSA*_ are available on github.com/3BioCompBio/EnzymeStability.

**Figure 1:**
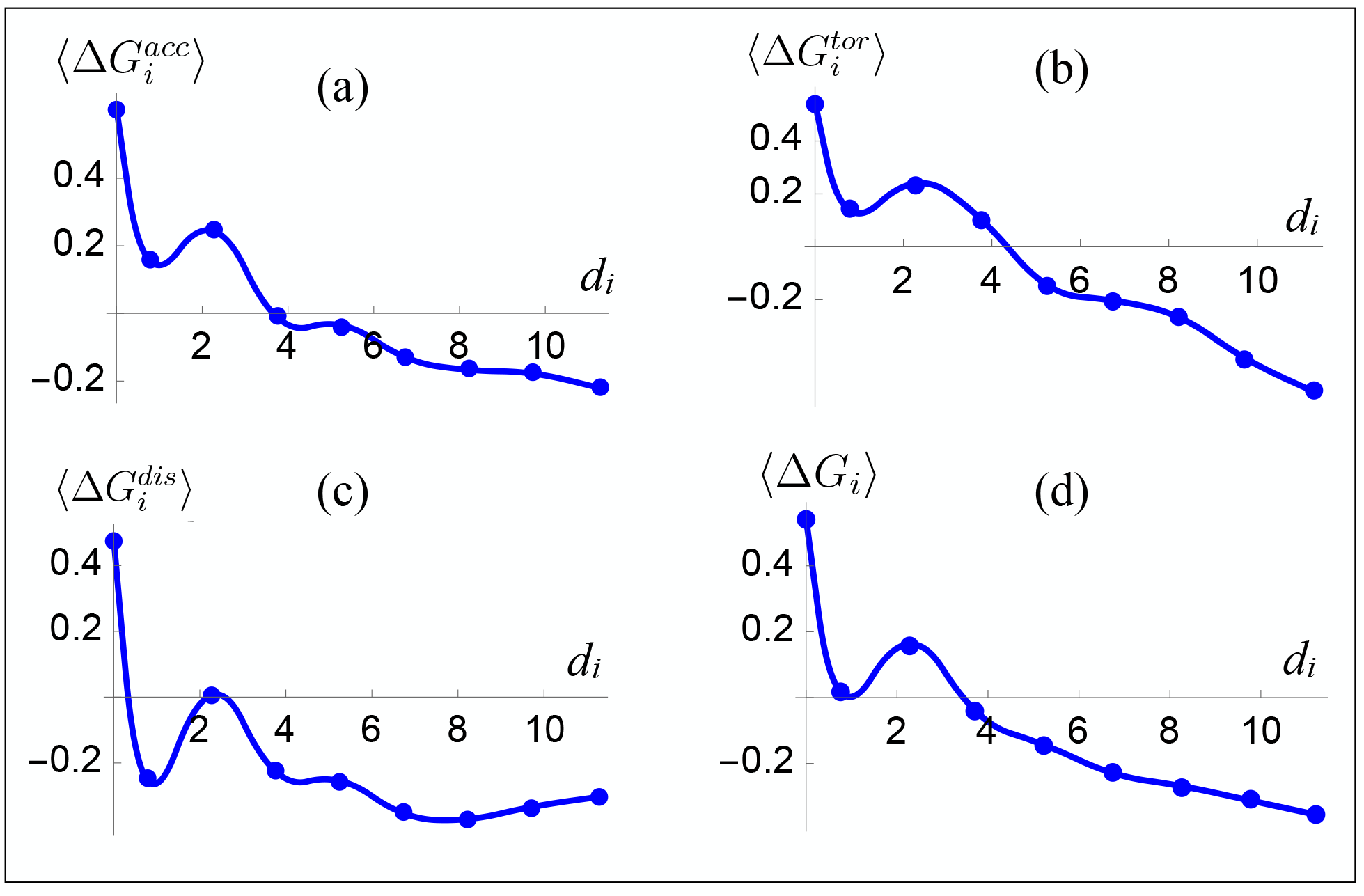
Overall folding free energy contribution ⟨Δ*G*_*i*_ ⟩ (in kcal/mol) of each residue *i* as a function of its distance *d*_*i*_ (in Å) from the closest catalytic residue. The folding free energy contributions were averaged over bins of 1.5 Å width and over all proteins from the 𝒟_*CSA*_ set. The blue dots are these average values and the curves were obtained by interpolation using Wolfram Mathematica [41]. The folding free energy contributions were computed with: (a) the solvent accessibility potential 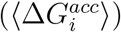; (b) the backbone torsion angle potential 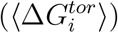; (c) the inter-residue distance potential 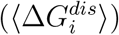 and (d) their combination (⟨Δ*G*_*i*_ ⟩).

**Figure 2:**
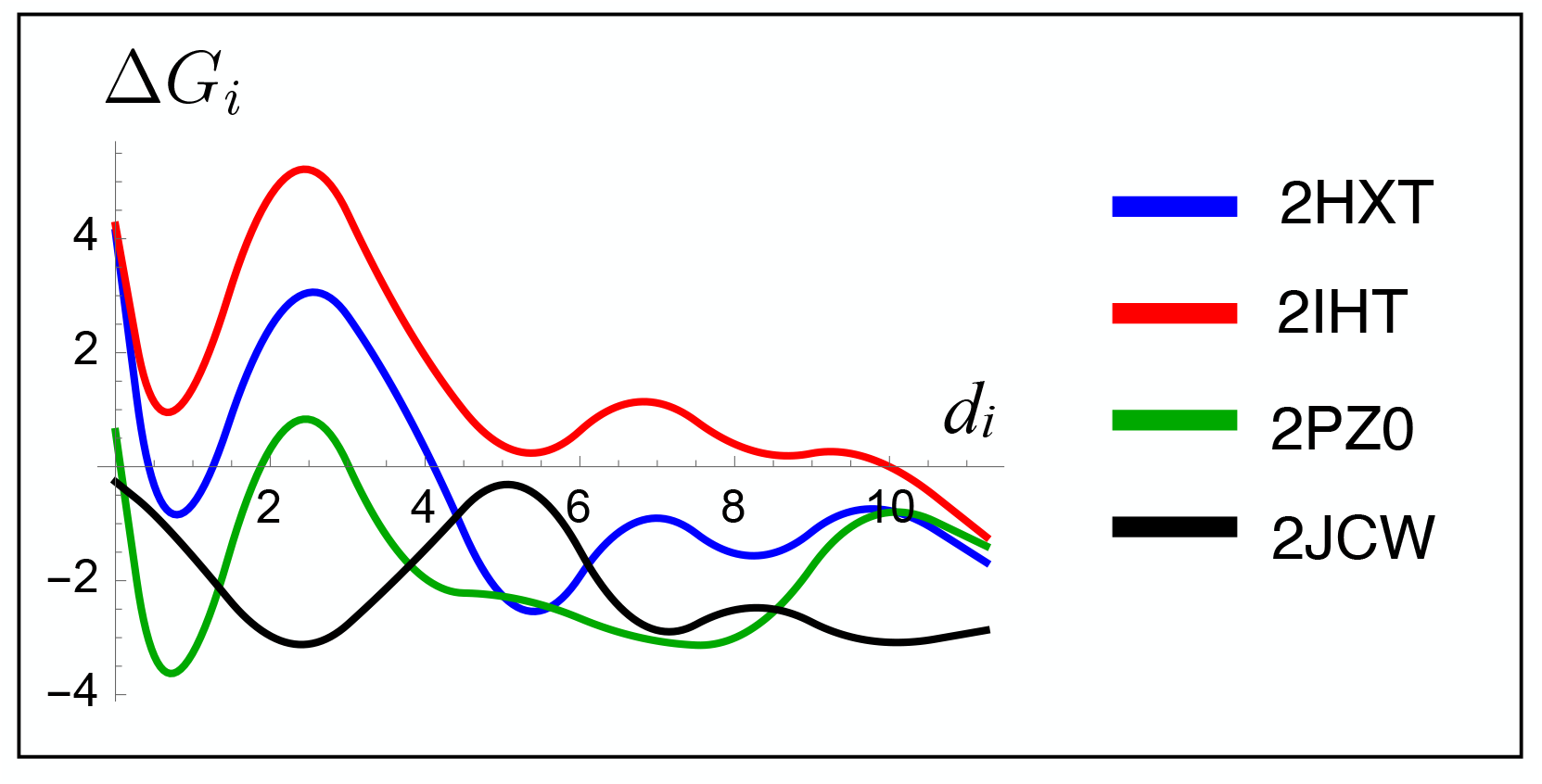
Per-enzyme folding free energy contribution ⟨Δ*G*_*i*_ ⟩ (in kcal/mol) of each residue *i* as a function of its distance *d*_*i*_ (in Å) from the closest catalytic residue, averaged over bins of 1.5 Å width. The curves were obtained by interpolation [41] of the averaged binned values. The enzymes considered are superoxide dismutase (PDB code 2JCW), carboxyethylarginine synthase (2IHT), L-fuconate dehydratase (2HXT) and aldo-keto reductase (2PZ0).

The first result we learn from Figs 1-2 is that catalytic residues are, on the average, stability weaknesses for all the statistical potentials considered. This is not surprising as they are optimized for their functional role rather than for stability, in agreement with previous reports [13, 18]. For example, the average Δ*G*_*i*_ values for all catalytic residues is equal to 0.54 kcal/mol, which is much higher than the average Δ*G*_*i*_ value of -0.37 kcal/mol obtained for all non-catalytic residues (Supplementary Table S1).

Interestingly, we observe a clear oscillatory pattern of the average per-residue folding free energy as a function of the distance to the catalytic site for most enzymes in 𝒟_*CSA*_ taken separately (Fig. 2 and github.com/3BioCompBio/EnzymeStability). The residues in the first shell around the catalytic site are more stabilizing, in the next shell they are weaker again, and so on. This effect is damped at larger distances, as the energetic compensation is more needed in the vicinity the active site to maintain the overall structure. The amplitude, period, vertical shift and damping of the oscillations differ according to the protein and, to a smaller extent, according to the potential (Supplementary Fig. S2).

In spite of this variability, an average oscillatory pattern emerges when considering all enzymes in 𝒟_*CSA*_ together, and this pattern is almost the same for all the potentials considered, as seen in Fig. 1. In the first shell, at about 1-2 Å from the catalytic center, the residues are less weak than the catalytic residues; they are sometimes even strengths. These residues still play a role in the catalytic reaction by energetically compensating for the important stability weaknesses of the catalytic residues. In the second shell, at distances of 2-3 Å, the residues become weak again, while at distances greater than 4 Å their contribution becomes increasingly stabilizing.

This stability compensation pattern appears as an emerging signature of enzymes in which residues close to the active site tend to counterbalance the stability weaknesses of the catalytic residues. Although the three considered potentials are defined from different conformational descriptors, they show a similar stability pattern, which gives further support to its generality. We would like to point out that the average oscillatory pattern of Fig. 1 appears despite the large variability of the per-enzyme folding free energy contributions visible in Fig. 2, with a standard deviation of about 0.5 kcal/mol.

One could argue that the observed pattern is the result of a non-trivial bias related to solvent accessibility, as residues in the core usually contribute more to stability than those at the surface. We thus repeated the same computation by averaging the Δ*G*_*i*_ values over all residues *i* that are situated either in the core (accessibility ≤ 15%) or at the surface (accessibility> 15%). The results plotted in Fig. 3 clearly show the same stability compensation patterns regardless of solvent accessibility. The only difference is the magnitude of the com-pensation, in other words, the height of the bump, which is about 0.2 kcal/mol in the core and less than 0.1 kcal/mol at the surface. The stability compensation in the core is thus stronger than at the surface, on the average. Note that this could be partly due to the averaging of the per-protein oscillatory patterns.

**Figure 3:**
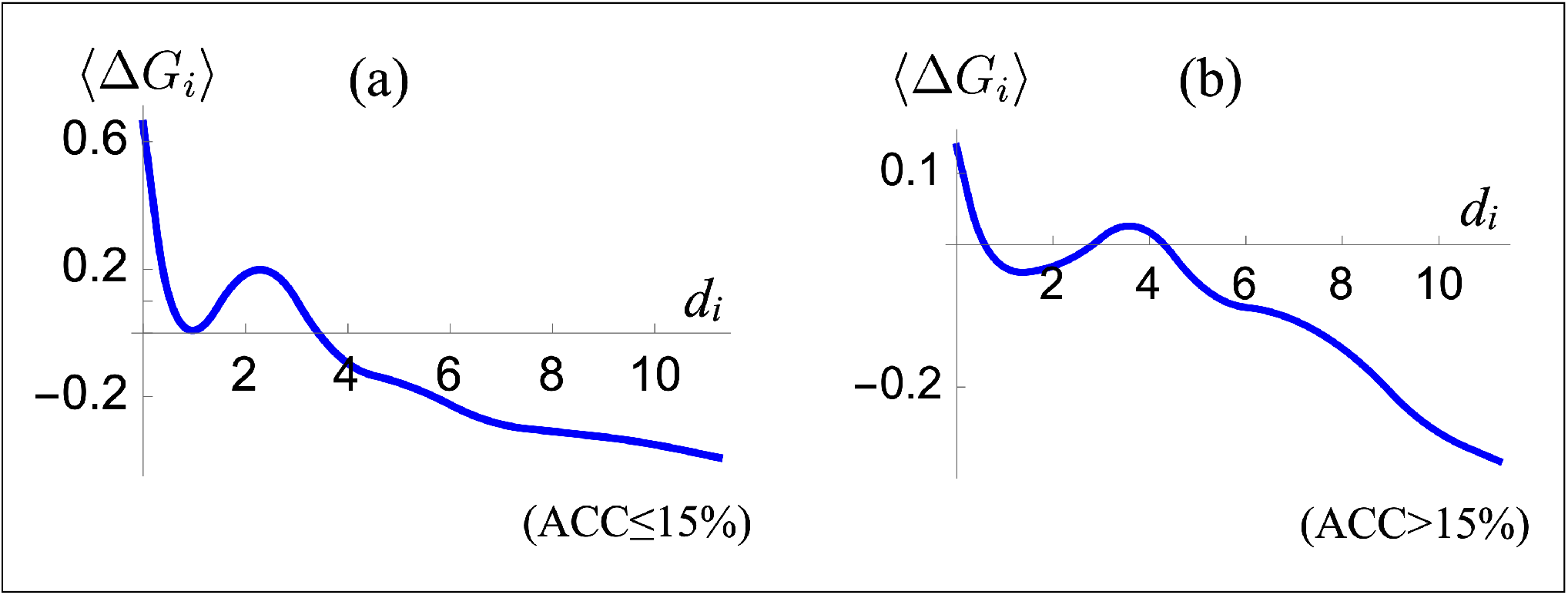
Folding free energy contribution ⟨Δ*G*_*i*_⟩ (in kcal/mol) for each residue *i* as a function of the distance *d*_*i*_ (in Å) from the closest catalytic residue. The folding free energy contributions were averaged over bins of 1.5 Å width and over all proteins from the 𝒟_*CSA*_ set. The curves were obtained by interpolation [41] of the averaged binned values. The residues *i* are limited to: (a) core residues with solvent accessibility smaller than or equal to 15%; (b) surface residues with solvent accessibility greater than 15%.

Finally, we investigated the possible differences in the observed strength/weakness patterns when focusing on different classes of enzymes. We considered for that purpose the enzyme commission (EC) nomenclature. We plotted in Fig. 4 the average ⟨Δ*G*_*i*_⟩ contributions as a function of the residue distance from the catalytic residues, separately for all enzymes belonging to the classes: EC1 (oxidoreductases), EC2 (transferases), EC3 (hydrolases), EC4 (lyases), EC5 (isomerases) and EC6 (ligases). The results of Fig. 4 clearly confirm that the stability compensation patterns are a universal trend shared by all enzymes, and are independent of the type of chemical reaction they catalyze. The strength of the compensation, however, slightly varies between enzyme classes.

**Figure 4:**
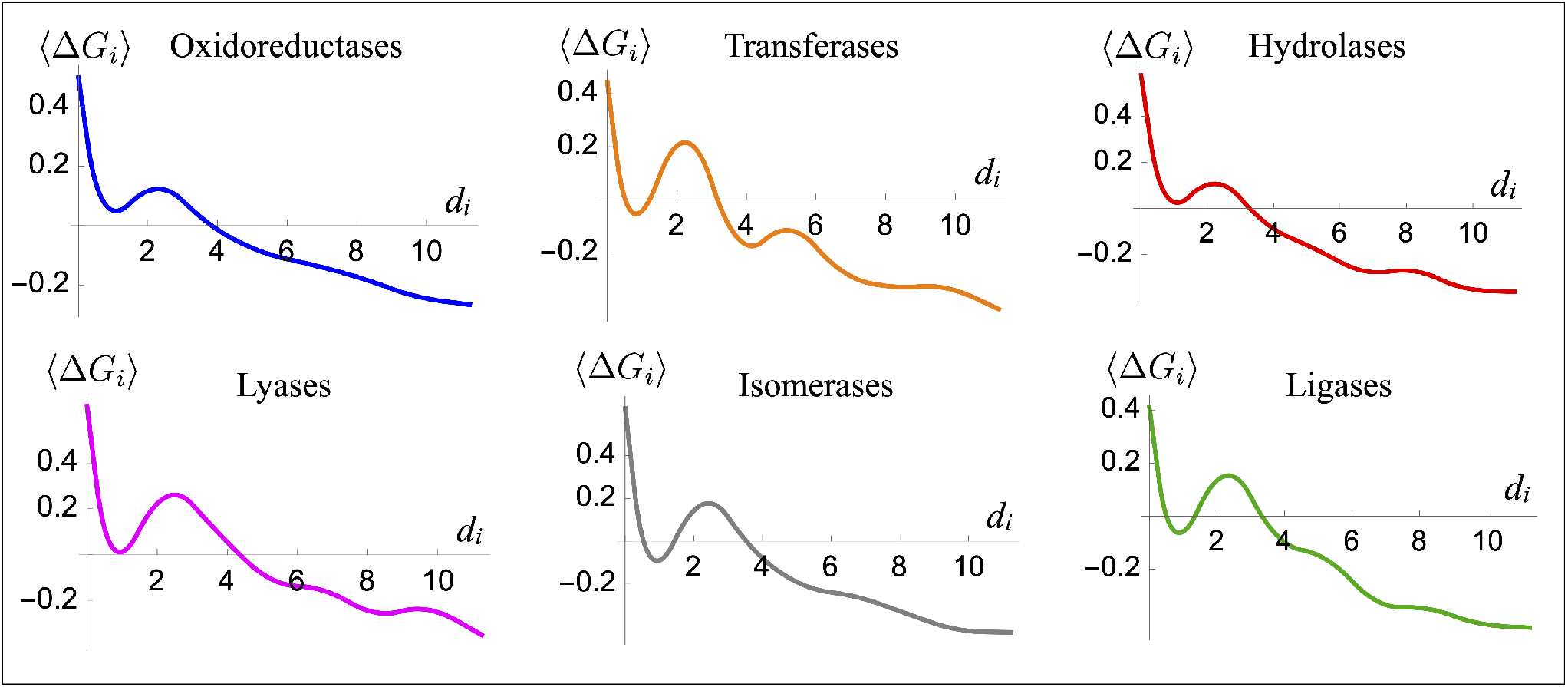
Folding free energy contribution ⟨Δ*G*_*i*_⟩ (in kcal/mol) of residue *i* as a function of its distance *d*_*i*_ (in Å) from the closest catalytic residue, averaged over bins of 1.5 Å width, for different EC classes of enzymes. The curves are obtained by interpolation [41] of the averaged binned values.

### 2.1 Strengths and weaknesses across evolution

Strength and weakness regions play a pivotal role in stability and function of proteins. It is therefore interesting to analyze how natural evolution shapes their interplay. Previous investigations have speculated that evolution tends to minimize the stability weaknesses of macromolecules or, in other words, their level of frustration [30, 17]. However, residual weaknesses are necessary for the enzymes to be functional [18]. To analyze the role of natural evolution, we estimated per-residue evolutionary rates using the ConSurf webserver [6], which generates and analyses multiple sequence alignments of protein families and outputs an evolutionary score 𝒮_*i*_ for each residue 𝒮*i*, with the convention that the lower _*i*_, the more conserved residue *i* in the protein family.

First of all, we plotted the average evolutionary score ⟨𝒮_*i*_⟩ as a function of residue distance *d*_*i*_ from the closest catalytic residue (Fig. 5.a). We also computed the Pearson correlation coefficient *r* between ⟨𝒮_*i*_⟩ and *d*_*i*_ and found a positive correlation with *r* equal to 0.39. This is expected, as catalytic residues are generally very well conserved for obvious functional reasons. But what is less expected is that there is a kind of non-trivial compensation similar to what we observed in the stability-distance plot of Fig. 1: catalytic residues are highly conserved; the closest residues in the 1-2 Å shell are a bit less conserved; the residues of the second shell, at distances of 2-3 Å from the catalytic residues, are again more conserved; and for the residues distant of more than 4 Å, conservation gradually decreases.

**Figure 5:**
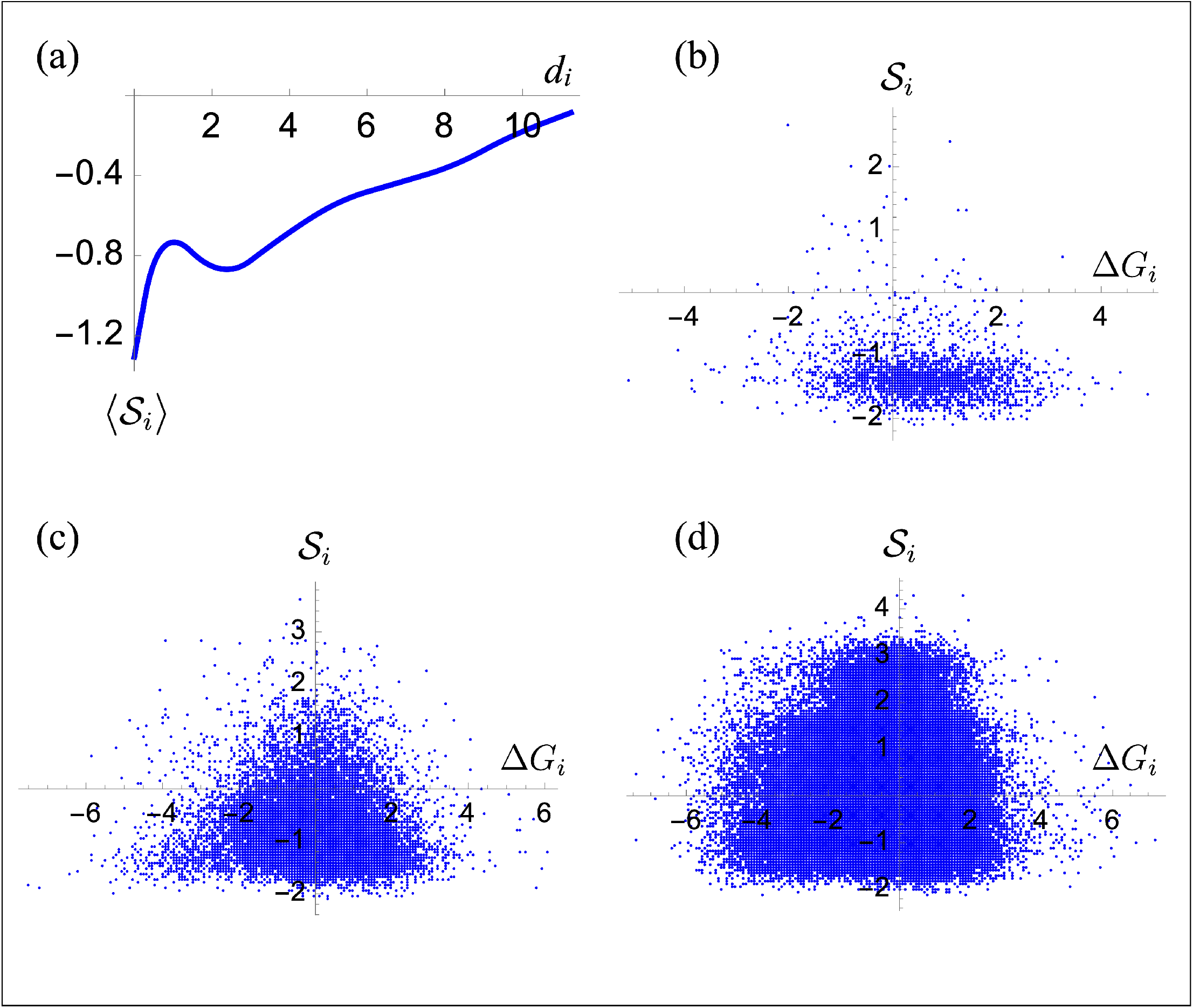
Relation between evolutionary conservation, folding free energy and distance to the active site. (a) Evolutionary score ⟨*S*_*i*_⟩ of residue *i* as a function of its distance *d*_*i*_ (in Å) from the closest catalytic residue, averaged over bins of 1.5 Å and all proteins from the 𝒟_*CSA*_ set. (b-d) Evolutionary score *S*_*i*_ as a function of the folding free energy Δ*G*_*i*_ (in kcal/mol) for (b) catalytic residues, (c) residues close to the catalytic site (0Å *< d*_*i*_ ≤ 5Å) and (d) all other residues.

To analyze this result in more detail, we examined the patterns that appear for individual enzymes. As shown in Supplementary Fig. S2, a clear oscillatory pattern emerges of ⟨𝒮_*i*_⟩ as a function of the residue distance *d*_*i*_, which is, however, much less pronounced than the Δ*G*_*i*_ oscillatory pattern. Interestingly, the oscillations are antiphased: shells of weaknesses correspond, on the average, to shells of well conserved residues, and shells of strengths, to shells of less conserved residues.

Although we found the distance from the catalytic site (*d*_*i*_) to be correlated with the residue conservation (𝒮_*i*_) and weakly anticorrelated with the folding free energy (Δ*G*_*i*_), we did not found any significant direct correlation or anticorrelation between 𝒮_*i*_ and Δ*G*_*i*_. Indeed, the Pearson correlation coefficient is equal to *r* = *−*0.03, and remains low when considering the three free energy contributions 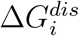, 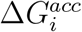 and 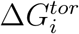 separately. This correlation is not surprising: a good anticorrelation would mean that all the weak residues are well conserved, and a good correlation, that all the strong residues are well conserved. Obviously, neither is true.

To explore this issue further, we plotted 𝒮_*i*_ as a function of Δ*G*_*i*_ for the whole 𝒟_*CSA*_ dataset containing more than 200k residues; we divided these residues into three groups: catalytic residues (Fig. 5b), residues in the periphery of the active site, i.e. at distances of 5 Å at most (Fig. 5c), and residues far from all active sites, at distances of more than 5 Å (Fig. 5d). We observe that catalytic residues are in general well conserved (𝒮_*i*_ *<* 0) and weak (Δ*G*_*i*_ *>* 0 kcal/mol). For the residues in the neighborhood of the catalytic sites, there are two categories: conserved residues (𝒮_*i*_ *<* 0) and poorly conserved residues (0 *<* 𝒮_*i*_ *<* 1.5); the former span a large range of Δ*G*_*i*_ values and can thus be either strong or weak, while the latter usually have neutral stability contributions (Δ*G*_*i*_ *≈* 0 kcal/mol). Among the residues far from the active site, many are well conserved (𝒮_*i*_ *<* 0) and many others not (𝒮_*i*_ *>* 0), many are weak (Δ*G*_*i*_ *>* 0 kcal/mol) and many others are strong (Δ*G*_*i*_ *<* 0 kcal/mol). But in general, there are more strong than weak residues in this category, and this is especially true for conserved residues (𝒮_*i*_ *<* 0 and Δ*G*_*i*_ *<* 0 kcal/mol).

We can thus conclude that, among the conserved residues, there is a substantial amount of weak residues and a substantial amount of strong residues. The former ensure the correct functioning of the enzyme and the latter, its overall structure and stability. This analysis clearly reflects the trade-off between stability and function and explains the insignificant correlation between folding free energy (Δ*G*_*i*_) and evolutionary conservation and (𝒮_*i*_).

### 2.2 Strengths and weaknesses as environmental adaptation mechanisms

We investigated if and how the patterns of stability strengths and weaknesses are tuned to adapt to different environmental conditions. It has long been known that cold-adapted proteins have a different enthalpy-entropy balance than heat-adapted proteins, so that they can maintain catalytic activity at low temperatures [1]. Multiple studies suggested that an improved flexibility in the regions around the active site is necessary to allow such enthalpy-entropy rearrangement [16, 31, 32, 14]. Recent computational analyses, however, point out that the whole surface outside the active site, which is usually not well conserved, is adapted through mutations in order to boost entropy contributions and achieve enzymatic rate enhancement at low temperature [2, 33].

To further investigate this issue, we first annotated all structures of the 𝒟_*CSA*_ dataset with the environmental temperature (T_*env*_) of their host organisms using data collected earlier [28]. We then formed two groups of proteins according to whether their host organisms are cold-adapted (T_*env*_ < 25°C) or heat-adapted (T_*env*_ ≥ 40°C). We computed the folding free energy value Δ*G*_*i*_ of all residues *i* in these two datasets as a function of their distance *d*_*i*_ to the active site; the results are shown in Fig 6.

**Figure 6:**
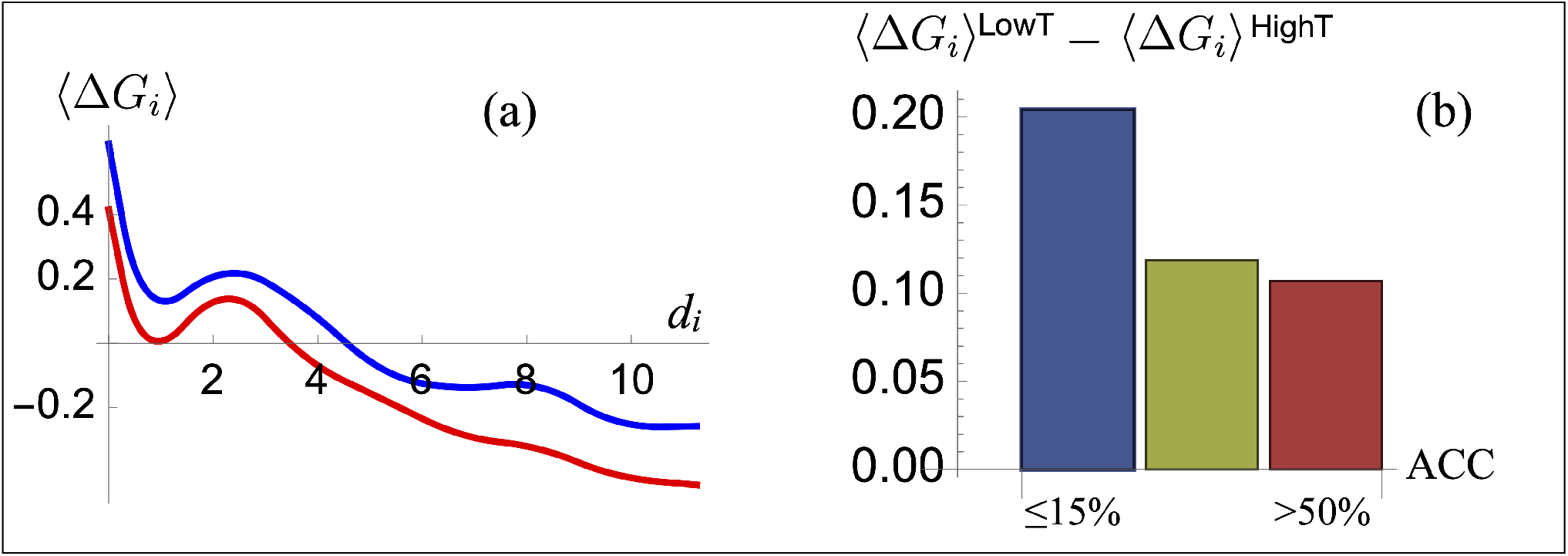
Difference in folding free energy between cold- and heat-adapted proteins from the 𝒟_*CSA*_ set. (a) Average per-residue folding free energy value ⟨Δ*G*_*i*_⟩ (in kcal/mol) as a function of the residue distance (in Å) to the closest catalytic site for residues in cold-adapted (blue) and heat-adapted proteins (red). (b) Difference in per-residue folding free energy (in kcal/mol) averaged over all residues *i* between cold- and heat-adapted proteins in three protein regions: core (accessibility *≤* 15%), partially buried (15% *<* accessibility *≤* 50%) and surface (accessibility *>* 50%).

We found that residues occurring in proteins from cold-adapted organisms are, on the average, weaker than residues from heat-adapted organisms. The average Δ*G*_*i*_ difference between the two sets is about 0.2 kcal/mol for any distance to the catalytic site (Fig. 6.a). Residues close to the catalytic site in cold-adapted organisms are thus weaker, as they have to allow an enhanced flexibility that in turn allows them to maintain the catalytic activity at low temperature.

We also analyzed the Δ*G*_*i*_ distribution of surface residues for the sets of cold- and heat-adapted enzymes, and found that the average Δ*G*_*i*_ of the former is higher by 0.1 kcal/mol than that of the latter. This trend is also observed in the partially buried region and in the core of the protein where it is even bigger, with a Δ*G*_*i*_ difference of 0.2 kcal/mol (Fig. 6.b). Our computation suggest that both flexibility in the active site regions and a general weakening of the structure play an essential role in the entropy-enthalpy shift of cold-adapted enzymes that allow them to remain functional in such lower temperatures.

### Comparison with experimental data

For comparing our computational results with experimental data, we used recently published fitness data obtained for the phosphatase and tensin homolog (PTEN) [25]. This enzyme is an oncosuppressor, which plays a fundamental role in the negative regulation of the proliferative phosphatidylinositol-3 kinase (PI3K) signalling pathway, by dephosphorylating the signaling lipid phosphatidylinositol (3,4,5)-trisphosphate (PIP3) [10, 24, 35]. Due to its involvement in cancer, lots of efforts have been devoted to understand the impact of mutations on its catalytic function.

We exploited deep mutagenesis scanning data, in which the *in vivo* impact of 7,244 single amino acid variants on the lipid phosphatase activity of PTEN was measured [25]. We computed the average of the measured fitness values of all single amino acid substitutions at a given residue position *i*, noted ℱ_*i*_, and compared it with the per-residue evolutionary conservation scores 𝒮_*i*_ computed using CONSURF [6] and with the structural stability values Δ*G*_*i*_ obtained using SWOTein [19]. We also computed the distance *d*_*i*_ of every residue *i* in PTEN to the closest of the active site residues Cys124 and Arg130.

Evolution (𝒮_*i*_) and fitness (ℱ_*i*_) values are, as expected, highly correlated with a Pearson correlation coefficient of 0.60 (P-value < 10^*−*31^). Conserved residues are indeed those that most contribute to protein fitness. Instead, stability (Δ*G*_*i*_) values do not shown a statistically significant anticorrelation with fitness (P-value = 0.52). This is illustrated in Fig. 7, where we observe their different behaviors when plotted as a function of *d*_*i*_. Note that if we limit ourselves to residues far from the catalytic center (*d*_*i*_ > 15 Å), Δ*G*_*i*_ and ℱ_*i*_ show a good anticorrelation with a Pearson coefficient of -0.26 (P-value < 0.001). Protein stability is thus an important component of fitness outside the catalytic site region: the stronger the residues, the higher their contribution to protein fitness, which results in a good ℱ_*i*_-Δ*G*_*i*_ anticorrelation. In the catalytic region, on the contrary, stability weaknesses and strengths both play a functional role and contribute to protein fitness; this explains the insignificant ℱ_*i*_-Δ*G*_*i*_ anticorrelation.

**Figure 7:**
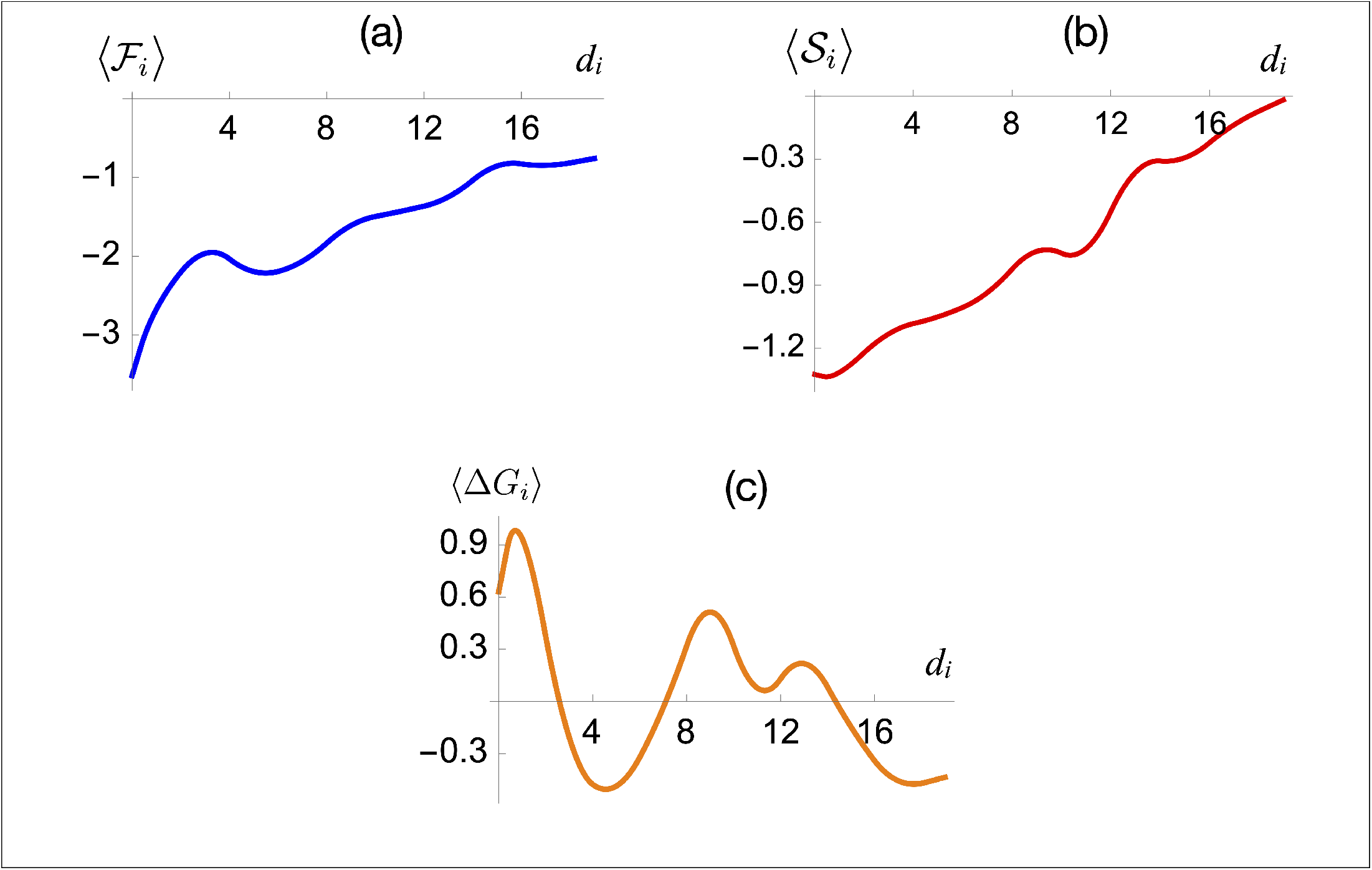
Fitness, stability and conservation properties in PTEN. (a) Measured per-residue fitness values ⟨ℱ_*i*_ ⟩ [25] as a function of the distance *d*_*i*_ (in Å) from the catalytic site averaged over 1.5 Å bins. (b) Residue conservation ⟨𝒮_*i*_ ⟩ as a function of *d*_*i*_ (in Å) averaged over 1.5 Å bins; (c) Per-residue folding free energy contributions ⟨Δ*G*_*i*_⟩ (in kcal/mol) as a function of *d*_*i*_ (in Å) averaged over 1.5 Å bins. The curves were obtained by interpolation [41] of the average binned values.

We continued by analyzing the correlation between the residue distance *d*_*i*_ to the active site and the per-residue fitness score ℱ_*i*_, conservation score 𝒮_*i*_ and stability value Δ*G*_*i*_. As expected, both fitness ℱ_*i*_ and conservation 𝒮_*i*_ correlate positively with the distance *d*_*i*_ with a Pearson correlation coefficient of 0.50 (P-value < 10^*−*21^) and 0.69 (P-value < 10^*−*44^), respectively. The closer residues are to the active site, the more their mutation has a negative impact on fitness. Consequently, variants close to the active site are more conserved during evolution than variants far from it.

The stability score Δ*G*_*i*_ is anticorrelated with the distance *d*_*i*_, as catalytic residues and residues in the vicinity are usually enriched in stability weaknesses. The correlation coefficient is weaker in absolute value than for fitness and evolution: -0.18 (P-value < 0.005). This is due to the non-linear, oscillatory, behavior of this score as a function of the distance, as we pointed out in the previous subsection. We can clearly observe this oscillatory pattern in Fig. 7: there are compensations between the stability weakness of the active site region, the stability strength of the first shell of residues at about 4-5 Å, and the stability weakness of the second shell of residues at about 8-13 Å. Note that this stability weakness-strength compensation is even more complex, as specific compensation patterns also occur within the different residue shells, as shown in Supplementary Fig. S3.

Mutations inserted close to the catalytic region have in general a negative impact on protein activity, but this is not the case for mutations further away from this region. It is therefore of interest to identify and analyze the positions in PTEN whose mutations lead to an increased catalytic activity. Here we focused on the ten positions of which the mutations most increase protein activity to find some share charateristics of these mutations. We first noticed that they are all situated on the protein surface, generally far from the catalytic area, at an average distance of about *d*_*i*_ *∼* 13 Å. Two main characteristics are shared by these positions: the wild-type amino acids are not conserved (⟨ 𝒮_*i*_⟩ = 0.2) and correspond to a stability weakness as their averaged Δ*G*_*i*_ is 0.6 kcal/mol. This result suggests that targeting non-conserved regions that are far from the catalytic area and that are stability weaknesses could be an interesting strategy to increase enzyme activity.

## 3 Discussion

The precise understanding of the trade-off between the two key properties of enzymes, activity and stability, and of how this trade-off is shaped by natural evolution, is important for both theoretical and practical reasons. In particular, it would help design or redesign enzymes that are optimized to work in conditions that differ from the physiological conditions. In this paper we applied our previously developed algorithm, SWOTein [19], to gain insights into this fundamental issue. We summarize our main results below.

First of all, we found a hallmark of the stability in the catalytic region, consisting of an energetic compensation between the catalytic residues which are usually stability weaknesses and their neighboring residues which are rather stability strengths or at least much less weak. A second shell of weak residues is located at distances of 2-3 Å from the catalytic center, these shells are surrounded by damped oscillatory patterns of weaknesses and strengths. This hallmark is general for all types of enzymes, *i*.*e* it does not depend on the catalytic reaction that they catalyze and does not depend on the solvent accessibility of the residues. Moreover, we showed that very similar stability patterns are observed for the three potentials 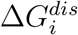, 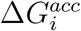and 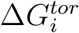 describing different types of residue interactions. This highlights that the stability compensation patterns involve not only tertiary interactions and hydrophobic forces, but also local interactions along the polypeptide chain.

Our results are in general agreement with an earlier study on frustration in catalytic regions [18], where a slight compensatory behavior between frustrated and less frustrated residue-residue interactions was observed. Stability strengths/weaknesses and frustration are related concepts: the former estimates the favorable or unfavorable contribution of each residue to the stability of the overall fold, and the latter compares the strengths of wild-type and mutated residue-residue interactions.

We also found that residue conservation across evolution shows a non-trivial compensation pattern that is similar to, but less marked than, the weakness and strength pattern: the catalytic residues are essentially conserved and weak; residues in the 1-2 Å shell around the catalytic center are less conserved and less weak; residues in the 2-3 Å shell are again more conserved and more weak; at distances of more than 4 Å the residue conservation continues to decrease and the residues become more and more stability strengths. In contrast, a linear relationship between evolutionary conservation and distance to the nearest catalytic residue was found in a earlier study [21], without any compensatory behavior. The disagreement between their results and ours is likely due to their use of wider shells of 5 Å, within which compensation takes place, whereas we considered thinner bins of 1.5 Å.

Interestingly, although the conservation and stability patterns have antiphased behavior, they are only weakly correlated. This is due to the fact that evolutionary conservation is highly correlated with enzyme fitness, while stability is only one of its main ingredients, the other being function. Note that the weight of stability in fitness increases when moving away from the active site.

We also applied our approach to study the adaptation of enzymes to different temperature conditions. By comparing the stability patterns of cold-adapted and heat-adapted proteins, we found that the former are characterized by a global weakening of the protein structure. More precisely, we observed not only a weaker catalytic pocket but also weaker surface and core regions in cold-adapted proteins than in heat-adapted homologues.

Finally, our study has important perspectives in enzyme design. Indeed, we can take advantage of what we have learned about stability and evolution to identify positions to target in order to increase enzyme activity. These positions are often far from the catalytic site, poorly conserved and, above all, stability weaknesses. The development of an enzyme activity improvement pipeline will be the subject of a forthcoming study. We would like to emphasize that, currently, the only way to identify functional positions to mutate in order to modulate function requires computationally expensive quantum mechanics/molecular mechanics methods [20]

## Acknowledgement

We acknowledge financial support from the FNRS - Fund for Scientific Research of Belgium through a PDR Research Project; MR is FNRS Research Director.

