## Supplementary for "Enzyme stability-activity trade-off : new insights from protein stability weaknesses and evolutionary conservation"

**Table of Content**

- Section S1. Statistical potentials and per-residue folding free energies.
- Table S1. Per-residue folding free energy contributions averaged on all core residues, surface residues, catalytic residues and on all protein residues.
- Figure S1. Examples of per-enzyme folding free energy contributions $\Delta G_i^{tor}$, $\Delta G_i^{dis}$ and $\Delta G_i^{acc}$ as a function of the distance to the closest catalytic residue.
- Figure S2. Folding free energy contribution $\Delta G_i$ and evolutionary conservation $\mathcal{S}_i$ of each residue as a function of its distance to the closest catalytic residue.
- Figure S3. $\Delta G_i$ values for all residues of PTEN that belong to a given distance range to the closest catalytic residue.

### Section S1. Statistical potentials and per-residue folding free energies

Statistical potentials are mean force potentials that have been widely applied to study protein stability. They are derived from the frequency of association between a structure element $c$ and a sequence element $s$ using the inverse Boltzmann law as [1, 2]:

$$\Delta G(c, s) = -k\,T\,\log \frac{P(c, s)}{P(c)P(s)} \tag{1}$$

where $P(c)$, $P(s)$ and $P(c, s)$ are the observation frequencies of $c$, $s$ or their association $(c, s)$ in a dataset of known 3D protein structures, $k$ is the Boltzmann constant and $T$ the absolute temperature. Note that we considered here globular proteins only; the mean force potentials derived from membrane proteins are different [3]. In our framework [4], we used three types of structural descriptors $c$, i.e. the backbone torsion angle domains, the inter-residue distances and the per-residue solvent accessibility, and one sequence element $s$ which is the amino acid type. We defined three different potentials, each associated to one structural descriptor, which we refer to as *tor* (torsion), *dis* (inter-residue distance) and *acc* (solvent accessibility) potentials.

These potentials were employed by our algorithm SWOTein [4] to compute three per-residue folding free energy contributions: $\Delta G_i^{acc}$, $\Delta G_i^{dis}$ and $\Delta G_i^{tor}$, where $i$ labels the residue positions, by partitioning the folding free energy into per-residue contributions as shown in [4]. These energy terms were used to describe the contribution of each individual residue $i$ to the overall protein stability. Residues that contribute unfavorably to the folding free energy have been defined as stability weaknesses while residues that contribute very favorably to it are considered stability strengths. We refer the reader to [4] for more details about the construction of our algorithm and on the choice of the parameters.

Here we briefly review some applications of our SWOTein method. In [5], we applied a first version of our method to study the strength/weakness patterns in bovine seminal ribonuclease; we found weaknesses among the catalytic residues and different weakness/strength patterns according to the oligomeric state of the protein, i.e. according to whether it is in its monomeric, dimeric or swapped dimeric state. In [4], we used SWOTein to perform large-sacle analyses and applied it to a redesigned apocytochrome $b_{562}$, compared the predictions with experimental data from native-state hydrogen-exchange experiments that identified the weakness regions in this protein, and found excellent agreement. We also successfully related the strength and weakness regions predicted by SWOTein to the large conformational changes occurring in the lysine/arginine/ornithine-binding protein from *Salmonella typhimurium*, and to molecular recognition upon formation of the complex between Subtilisin Carlsberg from *Bacillus subtilis* and the protease inhibitor Eglin c from *Hirudo medicinalis* [4].

The SWOTein free energy-based approach has remarkable advantages with respect to other commonly used computational predictions of protein weaknesses. In particular, it is much faster than molecular dynamics simulations where flexibility-based weaknesses are estimated from atomic root mean square deviations along the trajectories [6]. Indeed, SWOTein makes it possible to scan entire proteomes in just a few hours while maintaining excellent accuracy. Moreover, it is slightly faster and more accurate than frustration-based weakness predictions [7], as we showed earlier [4].

In this article, we used SWOTein's free energy contributions to systematically identify stability strength and weakness patterns in a set of 551 enzymes with known 3D structure and catalytic residues. We report in Table S1 the values of the per-residue folding free energy contributions computed with the three potentials, $\Delta G_i^{acc}$, $\Delta G_i^{dis}$ and $\Delta G_i^{tor}$, when averaged over all residues in the protein core, at the protein surface, at catalytic sites and over all residues in our dataset.

We first clearly observe that catalytic residues are, on the average, stability weaknesses of the protein structure. Indeed, they are optimized for functional rather than stability reasons, as discussed in the main text. Note the absence of major differences between the average values of the three free energy contributions for these residues: they are identified as weaknesses by all potentials.

Furthermore, we see that the free energy contributions have different behaviors in the protein core and at the surface. More precisely, the per-residue solvent accessibility folding free energy $\Delta G_i^{acc}$ has almost the same value when averaged over the core residues or the surface residues; the per-residue torsion folding free energy $\Delta G_i^{tor}$ has a more stabilizing effect at the protein surface, while the per-residue distance folding free energy $\Delta G_i^{sds}$ has a dominating stabilizing effect in the protein core as its main contribution is the hydrophobic effect [8]. As a metric of the per-site stability we chose to take $\langle \Delta G_i \rangle$ that is simply the average of these three contributions.

|  | Core | Surface | Catalytic | All |
|---|---|---|---|---|
| $\langle \Delta G_i^{acc} \rangle$ | -0.16 | -0.27 | 0.61 | -0.21 |
| $\langle \Delta G_i^{tor} \rangle$ | -0.36 | -0.98 | 0.54 | -0.62 |
| $\langle \Delta G_i^{dis} \rangle$ | -0.49 | 0.08 | 0.48 | -0.23 |
| $\langle \Delta G_i \rangle$ | -0.34 | -0.38 | 0.54 | -0.35 |

Table S1. Per-residue folding free energy contributions averaged on all core residues, surface residues, catalytic residues and on all protein residues. All values are in kcal/mol.

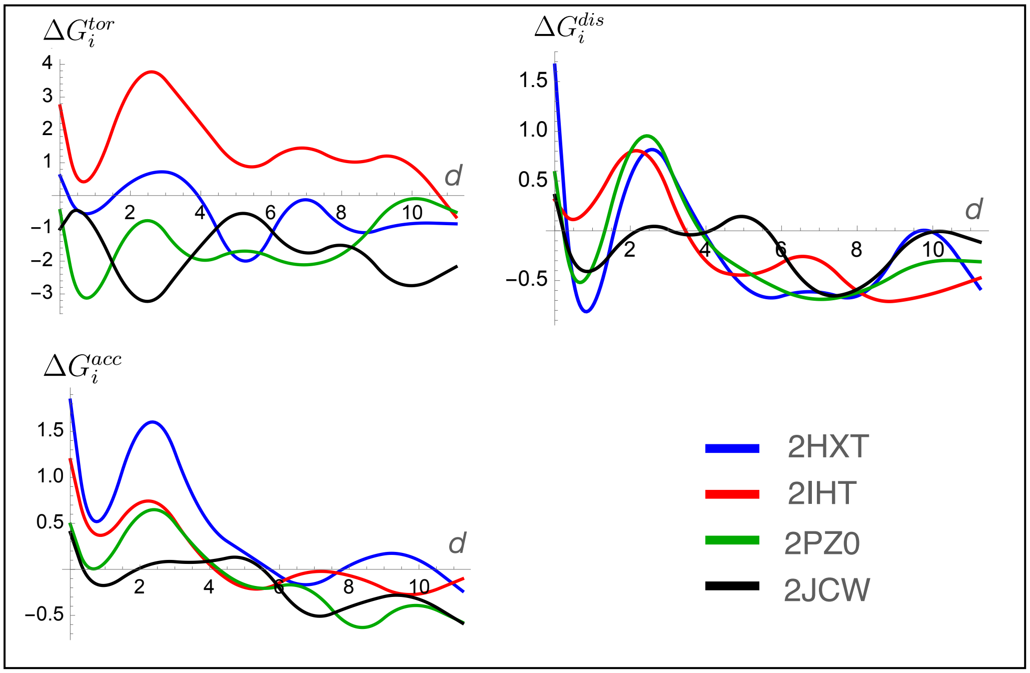


Figure S1. Examples of per-enzyme folding free energy contributions $\Delta G_i^{tor}$, $\Delta G_i^{dis}$ and $\Delta G_i^{acc}$ (in kcal/mol) of each residue $i$ as a function of its distance $d_i$ (in Å) from the closest catalytic residue, averaged over bins of 1.5 Å width. The curves were obtained by interpolation of the averaged binned values. The enzymes considered are superoxide dismutase (PDB code 2JCW), carboxyethylarginine synthase (2IHT), L-fuconate dehydratase (2HXT) and aldo-keto reductase (2PZ0).

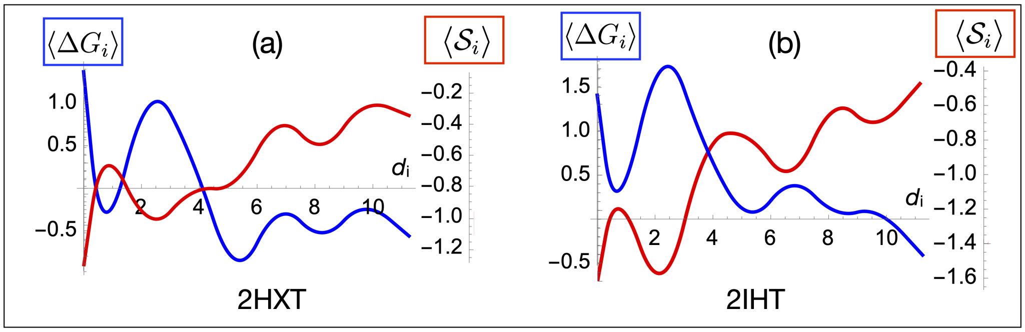


Figure S2. Folding free energy contribution $\Delta G_i$ (in kcal/mol) and evolutionary conservation $\mathcal{S}_i$ of each residue $i$ as a function of its distance $d_i$ (in Å) to the closest catalytic residue, averaged over bins of 1.5 Å width (blue and red curves, respectively). The curves were obtained by interpolation of the averaged binned values. The enzymes considered are carboxyethylarginine synthase (2IHT) and L-fuconate dehydratase (2HXT).

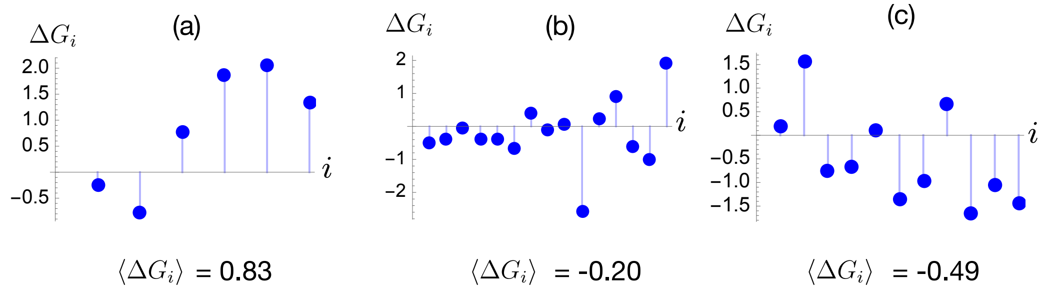


Figure S3. $\Delta G_i$ values (in kcal/mol) for all residues $i$ of PTEN that belong to a given distance range to the closest catalytic residue: (a) $0\text{Å} \leq d_i < 2\text{Å}$, (b) $2\text{Å} \leq d_i < 4\text{Å}$ and (c) $d_i \geq 4$ Å. The residues $i$ are given in sequential order but are not necessarily successive along the sequence. The average values $\langle \Delta G_i \rangle$ over all residues in each distance range is given below the plots.

[7] R Gonzalo Parra, Nicholas P Schafer, Leandro G Radusky, Min-Yeh Tsai, A Brenda Guzovsky, Peter G Wolynes, and Diego U Ferreiro. Protein frustratometer 2: a tool to localize energetic frustration in protein molecules, now with electrostatics. *Nucleic acids research*, 44(W1):W356–W360, 2016.

[8] Dimitri Gilis and Marianne Rooman. Predicting protein stability changes upon mutation using database-derived potentials: solvent accessibility determines the importance of local versus non-local interactions along the sequence. *Journal of molecular biology*, 272(2):276–290, 1997.
